# Exploring Disease Model Mouse Using Knowledge Graphs: Combining Gene Expression, Orthology, and Disease Datasets

**DOI:** 10.1101/2023.08.30.555283

**Authors:** Tatsuya Kushida, Tarcisio Mendes de Farias, Ana C. Sima, Christophe Dessimoz, Hirokazu Chiba, Frederic B. Bastian, Hiroshi Masuya

## Abstract

**Background:** The RIKEN BRC develops and maintains the RIKEN BioResource MetaDatabase to help users explore appropriate target bioresources for their experiments and prepare precise and high-quality data infrastructures. The Swiss Institute of Bioinformatics develops two RDF datasets across multi species for the study of gene expression and orthology: Bgee and Orthologous MAtrix (OMA, an orthology database).

**Methods:** This study integrates the RIKEN BioResource knowledge graph with Resource Description Framework (RDF) datasets from Bgee, a gene expression database, the OMA, the DisGeNET, a human gene-disease association, Mouse Genome Informatics (MGI), UniProt, and four disease ontologies in the RIKEN BioResource MetaDatabase. Our aim is to explore which model organisms are most appropriate for specific medical science research applications, using SPARQL queries across the integrated datasets. More precisely, we attempt to explore disease-related genes, as well as anatomical parts where these genes are overexpressed and subsequently identify appropriate bioresource candidates available for specific disease research applications.

**Results:** We illustrate the above through two use cases targeting either Alzheimer’s disease or melanoma. We identified two Alzheimer’s disease-related genes that were overexpressed in the prefrontal cortex (APP and APOE) and 21 RIKEN bioresources predicted to be relevant to Alzheimer’s disease research. Furthermore, executing a transitive search for the Uberon terms by using the Property Paths function, we identified two melanoma-related genes (HRAS and PTEN), and eight anatomical parts in which these genes were overexpressed, such as the “skin of limb” as an example. Finally, we compared the performance of the federated SPARQL query via the remote Bgee SPARQL endpoint with the performance of a centralized SPARQL query using the Bgee dataset as part of the RIKEN BioResource MetaDatabase.

**Conclusions:** As a result, we demonstrated that the performance of the federated approach degraded. We concluded that we improved the degradation of the query performance of the federated approach from the BioResource MetaDatabase to the SIB by refining the transferred data through the subquery and enhancing the server specifications, that is optimizing the triple store query evaluation.

## 1. Introduction

Bioresources are biological materials used for experimental life science research. They are widely used to elucidate the mechanisms of biological processes, including functional analyses, drug discovery, breeding, and practical chemical compound production as examples. Researchers generally source their bioresources from dedicated centers worldwide. These bioresource centers must develop retrieval systems to help users explore appropriate target bioresources for their experiments and to prepare precise and high-quality data infrastructure.

The BioResource Research Center (BRC) at the Japanese Institute of Physical and Chemical Research (RIKEN) is one of the largest and most comprehensive resource centers and manages a wide array of bioresources, such as experimental mouse strains, cultured cell lines and genetic material of human and animal origin, plant seeds, and microorganisms. The mission of the BRC is to contribute to the improvement of living standards, and the development and prosperity of human beings through distribution of its bioresources. These bioresources are developed and prepared under rigid quality control, so as to provide reliable infrastructure to firmly underpin life science research development. For its bioresource data infrastructure, RIKEN BRC adopted the Resource Description Framework (RDF), due to its advantages for data interoperability and its current adoption by institutions of the BRC’s interest for reuse. RIKEN BRC is working to continuously provide high-quality information by developing metadata and knowledge graphs (KG) and providing information retrieval systems.

The RIKEN BRC develops and maintains the RIKEN BioResource MetaDatabase (MetaDB) [1, 2]. This database integrates RIKEN BioResource RDF data with several life science datasets to support researchers in making a comprehensive use of RIKEN BRC’s research results. We call the integrated bioresource data the “RIKEN Bioresource Knowledge Graph.” So far, we have integrated the KG with the Orthologous MAtrix (OMA) database [3], DisGeNET [4], and disease ontologies, including MONDO Disease Ontology [5], Human Disease Ontology (DOID) [6], Orphanet Rare Disease Ontology (ORDO) [7], and Nanbyo Disease Ontology (NANDO) [8], which are provided by external organizations (Fig. 1). As a result, we are able to fully explore RIKEN BRC experimental mice, cell lines, and genetic materials available for research purposes [9].

**Fig. 1.**
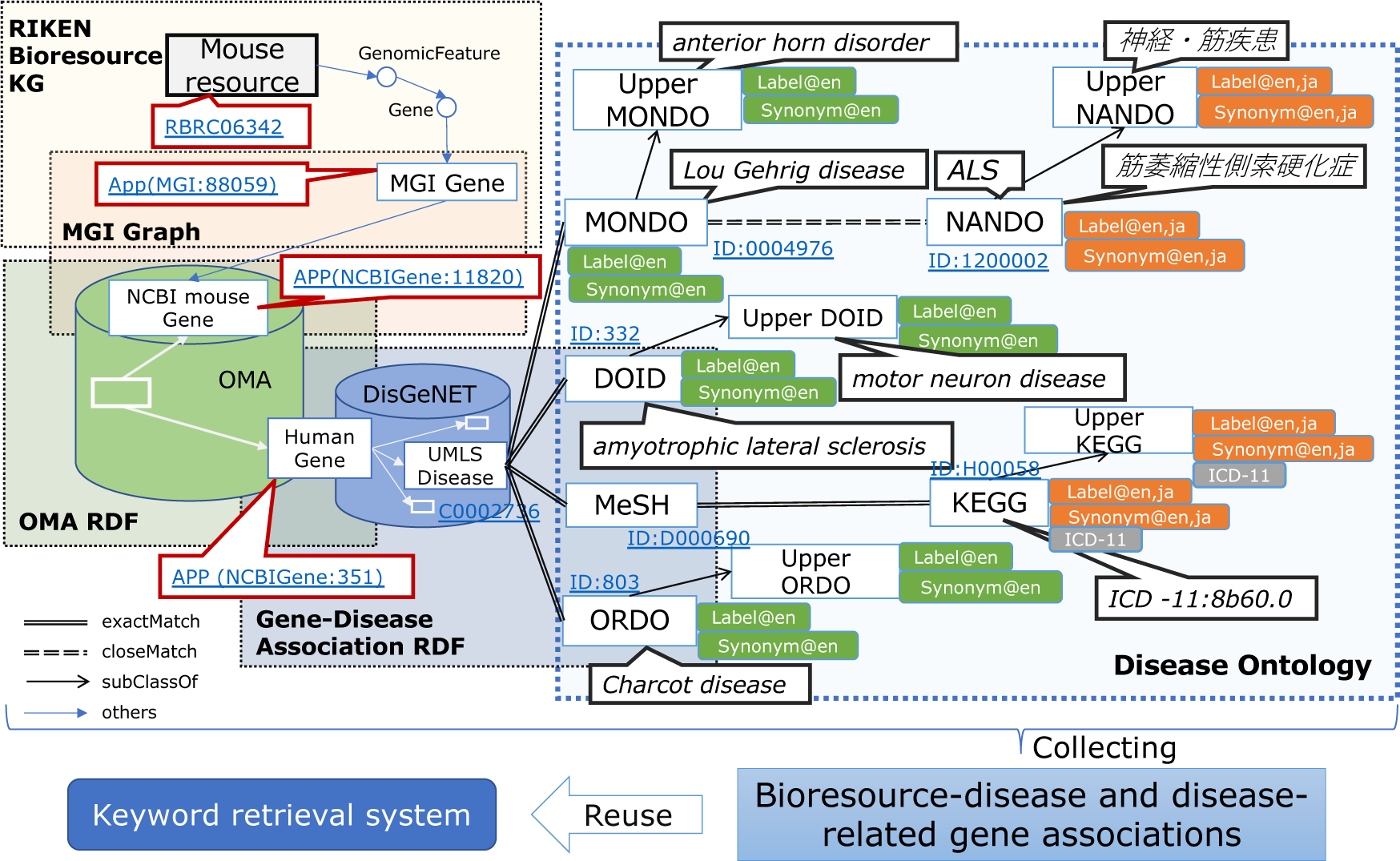
Data schema of RIKEN BioResource RDF data integrated with external RDF data and ontologies.

The SIB - Swiss Institute of Bioinformatics develops and maintains a growing catalog of publicly accessible knowledge graphs across many disciplines in the life sciences. For this study, we used two of the SIB RDF datasets for the study of gene expression and orthology: Bgee and OMA. Bgee [10] is a well-established gene expression database that integrates curated healthy wild-type expression data across a wide range of data sources to provide a comparable reference of normal gene expression across multiple animal species. OMA [3] (Orthologous MAtrix) is a database of orthologs among complete genomes across a wide range of species spanning the entire tree of life. Orthologs are pairs of genes that have evolved from a single gene in their last common ancestor. The OMA database provides orthologous information in the form of Hierarchical Orthologous Groups (HOGs), which are defined as gene families that contain genes that are all homologous to each other. The RDF version of OMA relies on the ORTH Ontology [11, 12].

In this article, we present a case study that combines the RIKEN BioResource MetaDB with Bgee for gene expression information, in addition to OMA for human-mouse orthologs and DisGeNET for associations between human genes and diseases (GDA). This case study investigates Alzheimer’s disease-(AD) and the melanoma-related human genes that are highly expressed in the healthy prefrontal cortex and the skin of body, respectively, and RIKEN’s genetically modified mice related to these genes. The rest of the paper is organized as follows: Section 2 reviews related work describing representative KG development cases and research in the life sciences. Section 3 explains the RIKEN Bioresource KG, and Section 4 presents external datasets and ontologies integrated with the RIKEN Bioresource KG. In Section 5 we performed SPARQL queries to retrieve finally bioresource candidates suited to a given disease research application. Section 6 presents the performance comparison results using the remote (federated) query over Bgee’s official SPARQL endpoint compared to using the local datasets (centralized) in BioResource MetaDB over a set of representative SPARQL query examples. Section 7 discusses the outcomes acquired using the integrated KG, a revealed issue, and a solution. Section 8 outlines future work.

## 2. Related work

The Monarch Initiative is an international consortium working to expand the use of genome information in biology and biomedical research. The Monarch Initiative publishes RDF data related to bioresources [13]. The published RDF data include relationships between mouse genes provided by Mouse Genome Informatics (MGI) [14] and related diseases and genome variation data. However, the Monarch initiative does not provide an official SPARQL endpoint, and users need to implement the triple store themselves to use the RDF data.

The Knowledge Graph Hub (KG-Hub) [15, 16] is a collection of biological and biomedical Knowledge Graphs, including their component data sources. It is provided by the Berkeley Bioinformatics Open-source Projects (BBOP) of the Lawrence Berkeley National Laboratory. KG-Hub tools comprise kghub-downloader, Koza (for data transformation), and KGX (Knowledge Graph Exchange), and KG-Hub uses these tools to transform data sources into standalone Biolink Model [17] compliant graphs. KG-Hub currently includes seven biomedical KG projects, including KG-COVID-19 [18] and KG-OBO [19]. The above-mentioned Monarch KG is also developed using these KG-Hub tools. KG-OBO translates the biological and biomedical ontologies on OBO Foundry [20] into graph nodes and edges. Ontology graphs translated by KG-OBO include Gene Ontology [21], ChEBI [22], and Uber-anatomy ontology (Uberon) [23].

Ubergraph [24] is an RDF triple store which provides a SPARQL query endpoint to an integrated suite of OBO ontologies, and includes precomputed inferred edges allowing logically complete queries over those ontologies for a subset of axioms in the Web Ontology Language (OWL) [25], and allows users to more efficiently access the integrated semantic knowledge graph. Ubergraph currently includes 39 OBO ontologies, including GO, ChEBI, Uberon, Cell Ontology (CL) [26], Mammalian Phenotype Ontology [27], and Human Phenotype Ontology [28].

## 3. RIKEN Bioresource Knowledge Graph

RIKEN BRC publishes metadata related to managed experimental animals, cell lines, genetic materials, experimental plants, and microorganism strains on its webpage [29]. Furthermore, the BRC is also developing RDF data and integrating bioresource metadata with external public datasets to enhance information and knowledge relevant to these bioresources. The Biological Resource Schema Ontology (BRSO) [30] is an RDF data model for various model organisms and the types, such as individual, cell, and DNA, which is largely developed by the Database Center for Life Science (DBCLS), RIKEN, and National Institute of Genetics (NIG). RIKEN BRC is developing bioresource RDF data based on the BRSO (Fig. 2) [2].

**Fig. 2.**
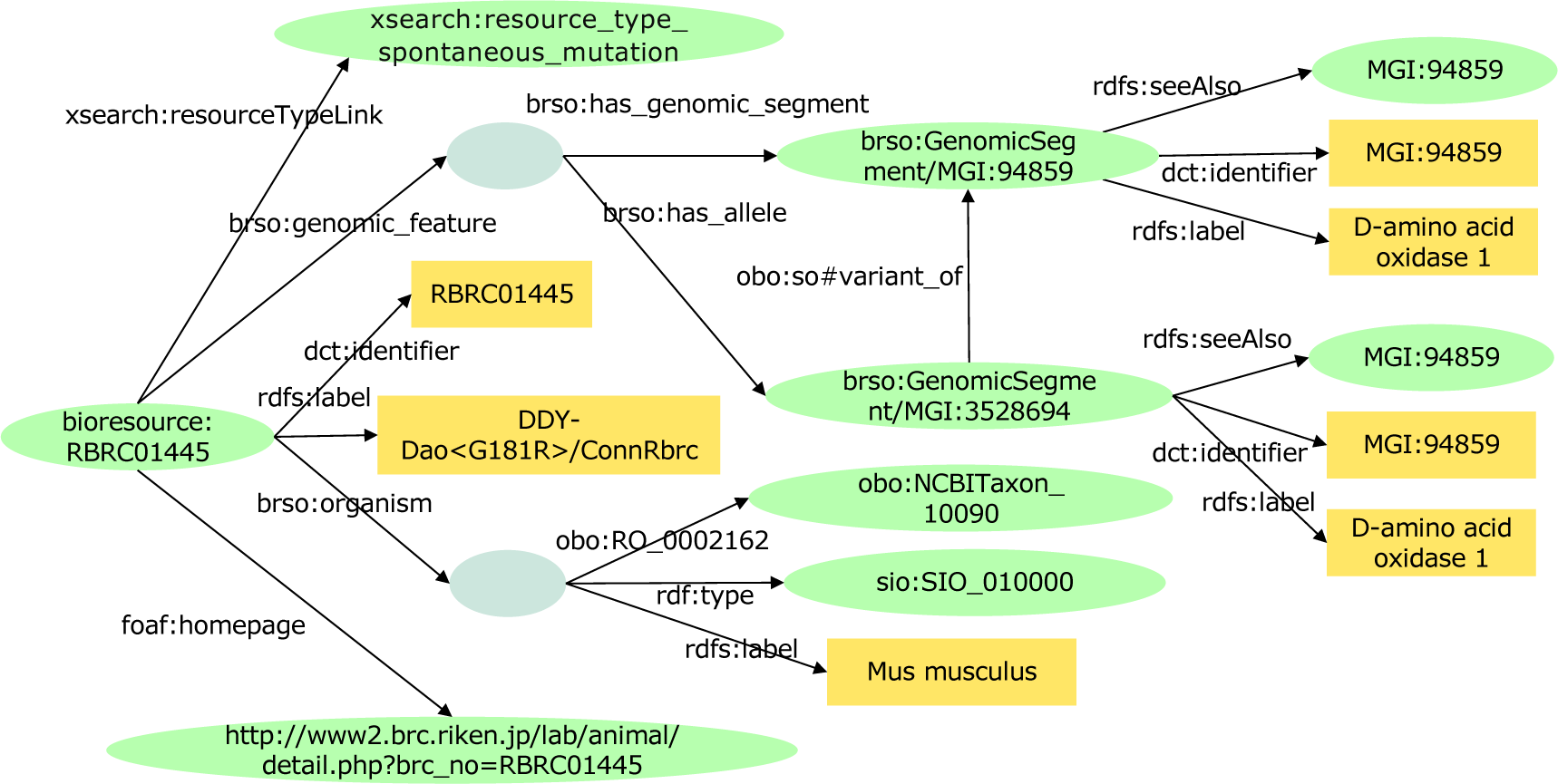
A part of RIKEN mouse (RBRC01445*) RDF data (KG) developed based on BRSO. *: https://knowledge.brc.riken.jp/resource/animal/card?lang=en&brc_no=RBRC01445

We term the RIKEN BRC bioresource RDF datasets the “RIKEN Bioresource Knowledge Graph.” The KG contains administrative information (e.g., bioresource developers, their affiliation), organisms (e.g., Mus muscles), bioresource types (e.g., spontaneous mutation mouse), gene id (e.g., MGI:94859), the related phenotypes and diseases (e.g., amyotrophic lateral sclerosis (ALS)). To date, we have developed KGs containing approximately 7,800 experimental mice, 9,600 cell lines, 125,000 genetic materials, 290,000 experimental plants, and 19,000 microorganisms. Users can browse KG data through a web interface, execute SPARQL queries, and download all the data from the BioResource MetaDB [31].

## 4. Data integration and interoperability

We are integrating the RIKEN Bioresource KG with external public datasets to enhance information and knowledge relevant to bioresources. Because almost all users are experimental researchers, the data retrieval system needs to enable researchers to explore candidate bioresources through a search of the KG using their familiar identifiers or keywords, such as MGI, NCBI, Ensembl Gene IDs and UniProtKB accession numbers. We therefore enhanced the KG to integrate the following information and knowledge:

### 4.1 MGI gene ID, Ensembl and NCBI gene ID mapping datasets

We developed RDF data representing relationships among MGI Gene ID, NCBI Gene ID, and Ensembl Gene ID from MGI Marker associations to Entrez Gene (tab-delimited) [32] provided from the MGI download page (Fig. 3). We stored the RDF data as a named graph in the BioResource MetaDB (Fig. 4). As a result, we could identify relationships between mouse resources, such as gene-modified mice and related NCBI Gene IDs and Ensembl Gene IDs in addition to MGI Gene IDs.

**Fig. 3.**
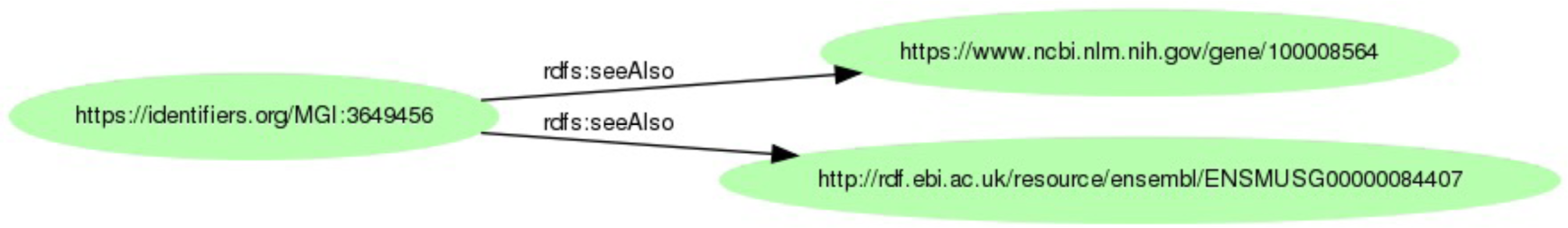
An example of RDF mapping data among MGI Gene, NCBI Gene and Ensembl Gene. This diagram was created using https://www.kanzaki.com/works/2009/pub/graph-draw.

**Fig. 4.**
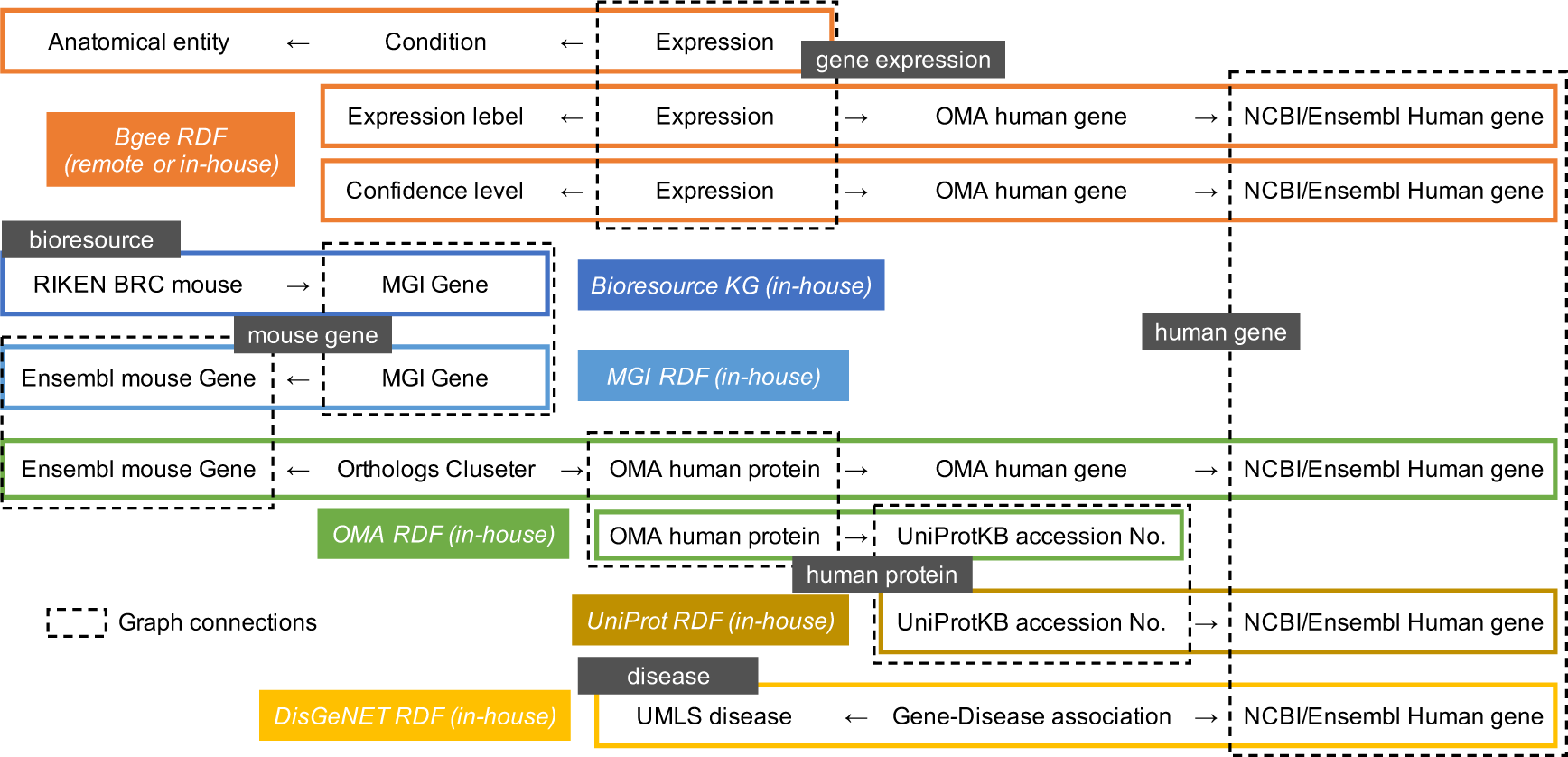
A simplified visualization of the query graph patterns.

### 4.2 UniProtKB accession number and NCBI gene ID mapping datasets

We developed RDF data representing relationships between the UniProtKB accession number and the NCBI Gene ID based on tab delimited files provided by UniProt [33]. We stored the RDF data as a named graph in the BioResource MetaDB (Fig. 4). As a result, we could identify relationships between mouse resources, such as gene-modified mice and related UniProtKB accession numbers, in addition to gene IDs.

### 4.3 OMA RDF datasets

We integrated the Bioresource KG with ortholog RDF datasets: OMA developed and provided by the Swiss Institute of Bioinformatics (SIB) as a named graph (Fig. 3, 4). This allowed us to acquire information on human Ensembl and NCBI gene IDs and UniProtKB accession numbers from gene-modified mouse gene IDs and UniProtKB accession numbers that are orthologous to human genes and proteins.

### 4.4 Bgee RDF datasets

We integrated the Bioresource KG with the gene expression RDF dataset Bgee, developed and provided by SIB as a named graph (Fig. 4). As a result, we could access information on gene expression patterns, confidence levels and expressed anatomical parts from human Ensembl and NCBI gene IDs and UniProtKB accession numbers.

### 4.5 Gene-Disease Association RDF datasets

We integrated the Bioresource KG with human gene-disease association RDF datasets: DisGeNET and MedGen as named graphs [34] (Fig. 3, 4). The former was developed by the Institute for Research in Biomedicine (IRB, Barcelona), and the latter was developed by National Center for Biotechnology Information (NCBI), and the RDF data were generated and provided by DBCLS. This study used the GDA datasets of which the GDA score was 0.5 or more extracted from DisGeNET RDF v7.0.0 in the RDF Portal [35, 36]. As a result, we could access information on related human disease identifiers, such as UMLS IDs or MedGen IDs (e.g., C0002736, amyotrophic lateral sclerosis (ALS)) from human Ensembl and NCBI gene IDs and UniProtKB accession numbers.

### 4.6 Disease Ontologies

We incorporated the OWL version of four disease ontologies that are used as controlled vocabularies: MONDO [5], DOID [6], ORDO [7], and NANDO [8] as named graphs, into the BioResource MetaDB (Fig. 3). The Monarch Initiative developed MONDO. The University of Maryland mainly developed DOID. ORDO was mainly developed by the National Institute of Health and Medical Research (INSERM) and the European Bioinformatics Institute (EBI). NANDO was mainly developed by DBCLS and RIKEN. As a result, we could access information on related human gene IDs from English and Japanese disease names, Disease Ontology IDs, and ICD-11 (International Classification of Diseases 11th Revision) [37] through these ontologies and DisGeNET.

## 5. Exploring bioresources relevant to human diseases

In this study, we aim to identify disease-related genes, the anatomical parts where the genes were overexpressed, and the RIKEN bioresource relevant to the disease, by exploring the extended Bioresource KG using SPARQL queries. We applied this to two concrete use cases, targeting the study of Alzheimer’s disease and melanoma.

**Example 1-1**: Federated query for Alzheimer’s disease (see Additional file 1) is a query for exploring AD-(UMLS:C0002395) related genes overexpressed in specific anatomical parts (e.g., prefrontal cortex) and the bioresources expected to be available for AD research. This study partially revised SPARQL queries used in our previous report [38] to improve the query performance and executed the revised queries in the SPARQL endpoint [39] of the RIKEN BioResource MetaDB. The executed query includes the query conditions: the prefrontal cortex (UBERON:0000451) as location of overexpression, a high confidence level for expression data, and the expression score greater than 99 where the maximum score is 100. For the sake of simplicity, we consider overexpressed genes when the expression score is above 99. The greater the expression score, the greater is the expression of a gene in a given condition. We used the DisGeNET as gene-disease association datasets with the GDA score [4] of 0.5 or more.

We present the query results in Table 1. We observed that the APP gene (ENSG:00000142192) and APOE gene (ENSG:00000130203) were AD-related genes and that RBRC06342 and RBRC03391 were RIKEN mouse resources expected to be of relevance for AD research. APP and APOE genes have previously been linked to experimental AD, as reported in [40, 41]. The query runtime, on average, was approximately 59 seconds (Table 2).

**Table 1.** Results of Example 1-1: Federated query for Alzheimer’s disease and Example 2-1: Centralized query for Alzheimer’s disease.

| Query approach | No. of mice | RIKEN Mouse IDs | No. of genes | Ensembl Gene IDs | Gene labels |
| --- | --- | --- | --- | --- | --- |
| Example 1-1:<br>Federated query for AD | 21 | RBRC06342 RBRC06343<br>RBRC06344 RBRC10041<br>RBRC10042 RBRC10700<br>RBRC10701 RBRC10043<br>RBRC11518 RBRC03298<br>RBRC03388 RBRC03389<br>RBRC03390 RBRC03391<br>RBRC03418 RBRC03419<br>RBRC03420 RBRC03421<br>RBRC03422 RBRC03423<br>RBRC00052 | 2 | ENSG00000130203<br>ENSG00000142192 | APOE<br>APP |
| Example 2-1:<br>Centralized query for AD |  |  |  |  |  |

**Table 2.**
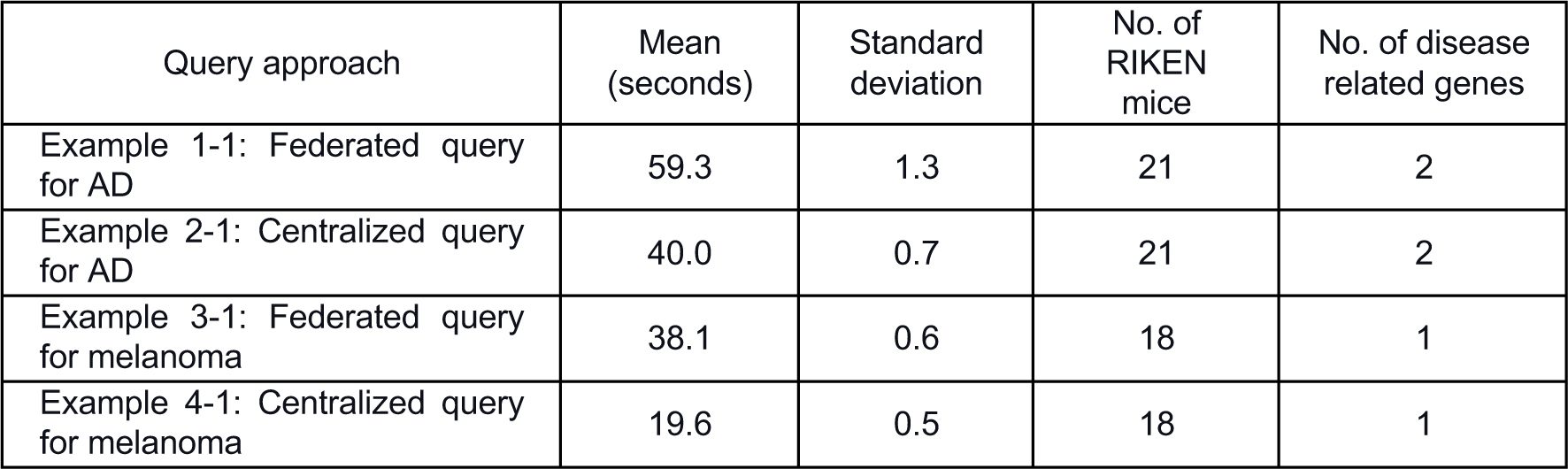
The query execution time of Examples 1-1, 2-1, 3-1, and 4-1. The queries were executed 10 times each at https://knowledge.brc.riken.jp/sparql.

| Query approach | Mean (seconds) | Standard deviation | No. of RIKEN mice | No. of disease related genes |
| --- | --- | --- | --- | --- |
| Example 1-1: Federated query for AD | 59.3 | 1.3 | 21 | 2 |
| Example 2-1: Centralized query for AD | 40.0 | 0.7 | 21 | 2 |
| Example 3-1: Federated query for melanoma | 38.1 | 0.6 | 18 | 1 |
| Example 4-1: Centralized query for melanoma | 19.6 | 0.5 | 18 | 1 |

In this study, we ran SPARQL query tests and obtained the retrieval results and runtimes on 4 August 2023.

## 6. Comparison between federated and centralized query performance

Furthermore, we evaluated two query execution scenarios [38]. One scenario considers a SERVICE SPARQL subquery to be executed against the remote Bgee SPARQL endpoint resulting, assuring access to the latest data. The second scenario replaces the centralized SPARQL query example with a subquery matching triple patterns from the named graph containing Bgee data and stored in the RIKEN BioResource MetaDB. We obtained the Bgee RDF data [42] on 20 July 2023 and incorporated it into the RIKEN BioResource MetaDB. To avoid longer runtimes and query timeout, we used the locally stored OMA and DisGeNET as named graphs in the BioResource MetaDB in both scenarios (Fig. 4).

**Example 2-1**: Centralized query for Alzheimer’s disease (see Additional file 2) is based on the second scenario. Table 1 shows the query results of examples 1-1 and 2-1. The results of both were identical. The average query runtime of Example 2-1 was approximately 40 seconds, and it was faster than that of Example 1-1 (Table 2).

We further compared federated versus centralized data access and storage approaches for other use cases. **Example 3-1** and **Example 4-1** are queries for melanoma (UMLS:C0025202) using the federated query (see Additional file 3) and centralized query (see Additional file 4) for Bgee data, respectively. These queries include the melanoma-related genes that were overexpressed in the skin of body (UBERON:0002097) as a query condition. The other query conditions were the same as the Examples 1-1 and 2-1.

Table 3 shows the query results of Examples 3-1 and 4-1. The findings were identical and included the demonstration that the HRAS gene (ENSG:00000174775) was overexpressed in the skin of body as a melanoma-related gene, and 18 RIKEN bioresources were expected to be relevant to the melanoma research, such as RBRC10866 [43] and RBRC01088 [44]. Table 2 shows the runtimes of Examples 3-1 and 4-1. The average runtime of Example 3-1 (using a federated query) was approximately 38 seconds, while that of Example 4-1 (using a centralized query) was approximately 20 seconds, about half of the time of Example 3-1.

**Table 3.**
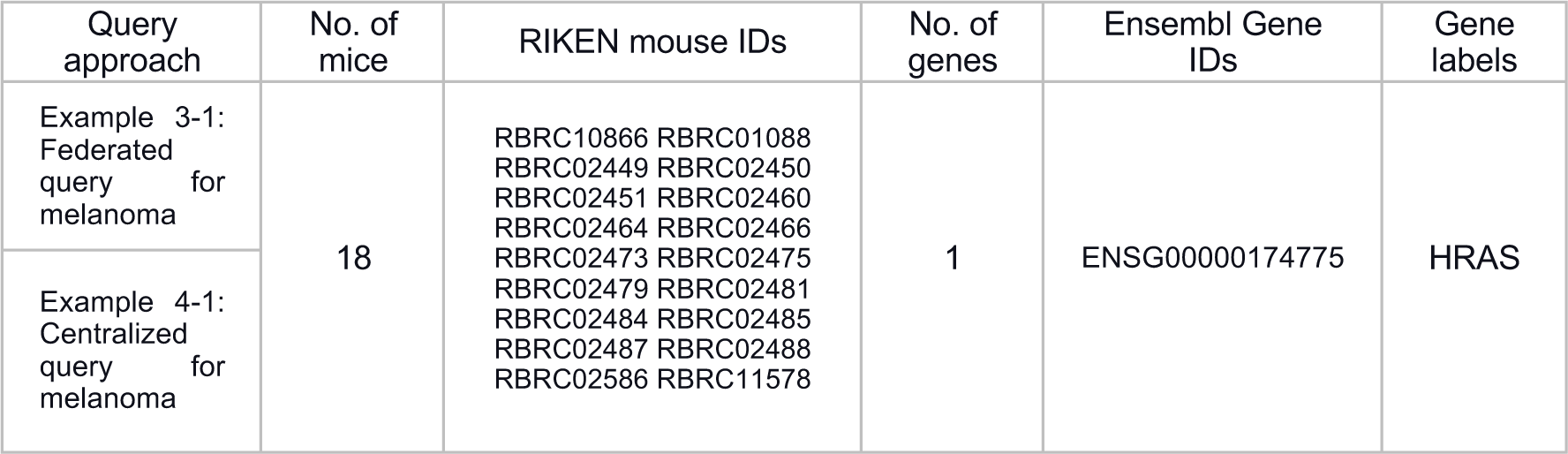
Results of Example 3-1: Federated query for melanoma and Example 4-1: Centralized query for melanoma.

In comparison between Examples 1-1 and 2-1, and 3-1 and 4-1, the query execution performance was significantly better in the centralized setup. Note that we executed the centralized queries Examples 2-1 and 4-1 for the same data as available via the remote Bgee SPARQL endpoint [45]. Thus, the significant differences between the federated and the centralized runtimes were not due to the Bgee data version. Mainly because in our experimental setup, we do not consider any cost-based federated query processing engines [46], and we expected that the performance of federated queries would be significantly worse than the same centralized query, notably, due to network latency and poorer query optimization plan of federated queries.

## 7. Discussion

### 7.1 Analysis and improvement of query performance

To ensure that bioresources are appropriately used as research materials in a wider range of studies, bioresource centers need to provide users with up-to-date and detailed information on the characteristics of bioresources. Integration of datasets for bioresources which are independently collected by bioresource centers with datasets from public databases maintained in biomedical databases is essential for this purpose. As a use case for the integration and exploitation of remote data using the Semantic Web technologies and RDF, we compared the performance of variations in the way SPARQL queries focused on subqueries and federated search.

Subqueries represent a way to embed queries within other SPARQL queries, normally to achieve results which cannot otherwise be achieved, such as limiting the number of results from some sub-expression within the query [47]. The appropriate usage of subqueries is expected to improve query performance. Examples 1-1, 2-1, 3-1, and 4-1 contain one subquery because we had not obtained the query results due to the transaction timeout in the case without using the subquery (data not shown). To estimate how the usage of the subquery will affect query performance, we divided the SPARQL queries into four sections and investigated how the arrangement of the subquery within the query could improve query performance (Fig. 5). **Examples 1-2** (see Additional file 5), **2-2** (see Additional file 6), **3-2** (see Additional file 7), and **4-2** (see Additional file 8) are queries with two subqueries, and the remaining query conditions are the same as Example 1-1, Example 2-1, Example 3-1, and Example 4-1. **Examples 1-3** (see Additional file 9), **2-3** (see Additional file 10), **3-3** (see Additional file 11), and **4-3** (see Additional file 12) are queries with three subqueries, and the remaining query conditions are the same as Examples 1-1, 2-1, 3-1, and 4-1, respectively. For example, in Example 1-2, Section 1 is nested inside Section 2. Furthermore, Sections 1, 2, and 3 are nested inside Section 4 (that is Bgee’s query). The nested query (subquery) is evaluated first, and the outer query uses the results.

**Fig. 5.**
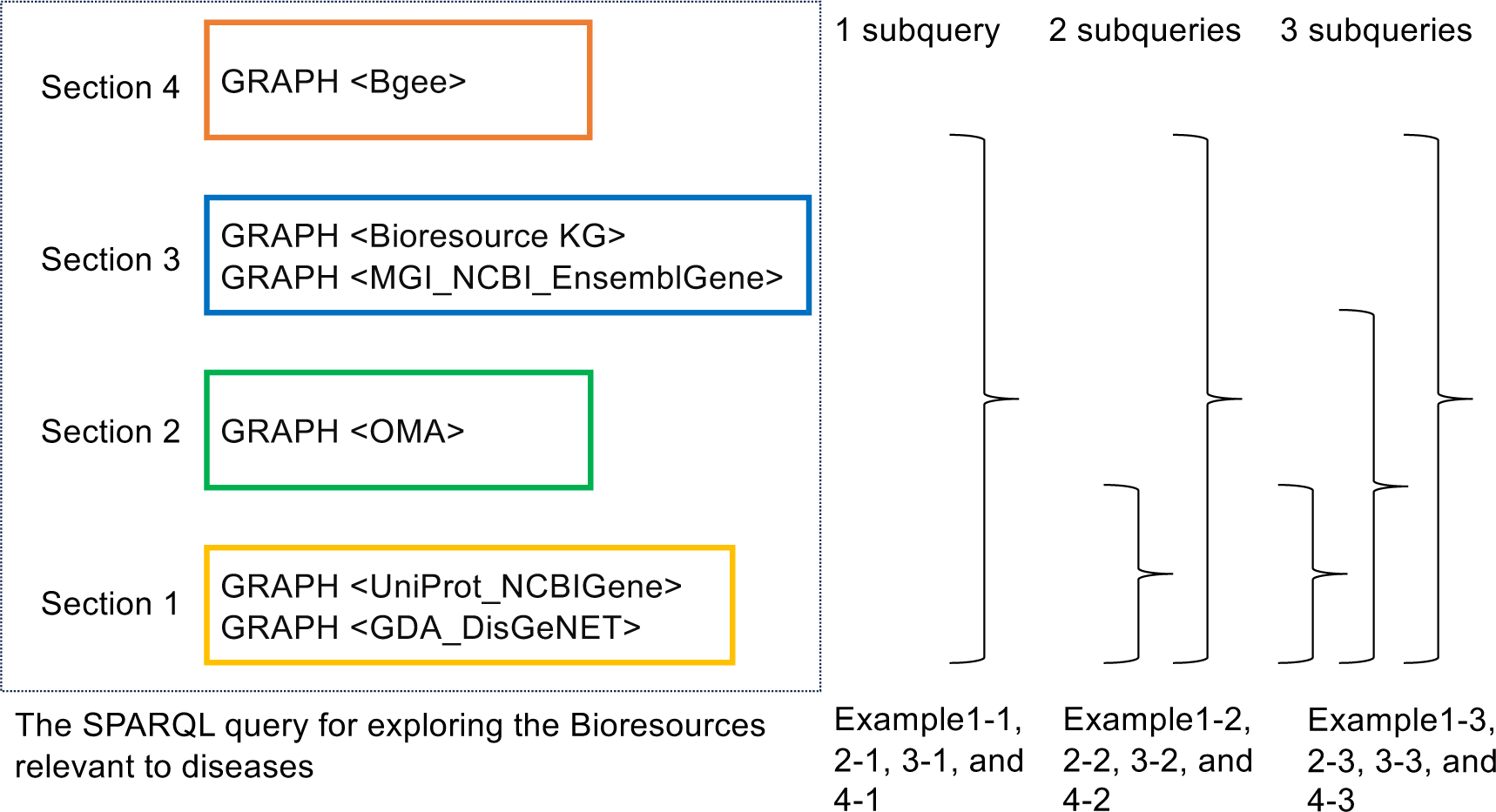
Four sections within the SPARQL query examples and the position of the subqueries. For the outline of query graph patterns, refer to Fig. 3.

Table 4 shows the average runtimes of Example 1-x, 2-x, 3-x, and 4-x. Bold values represent the best values in each column. For instance, in the row of Example 1-x (It means Examples 1-1, 1-2, and 1-3), Example 1-3 with three subqueries was shorter than the others. The most effective arrangement of the subqueries for the query performance was different in each row, although all example queries had the same graph structures. On the other hand, we observed that the performance of the query could be significantly improved when we used the subquery in particular places, thereby providing a more effective query plan. The differences between the average runtimes of Example 1-x (federated for AD) and Example 2-x (centralized for AD), and Example 3-x (federated for melanoma) and Example 4-x (centralized for melanoma) were about 25 and 7 seconds, respectively. The query results, such as the AD-related genes of Examples 1-2 and 1-3 were the same as those of Example 1-1, the results of Examples 2-2 and 2-3 were the same as those of Example 2-1 (Table 2), the results of Examples 3-2 and 3-3 were the same as those of Example 3-1, and the results of Examples 4-2 and 4-3 were the same as those of Example 4-1, respectively (Table 3). From these results, we confirmed the appropriate use of the subqueries for all Examples.

**Table 4.** The average runtime from 10 executions of the SPARQL query Examples 1-x, 2-x, 3-x, and 4-x, including one-time, twice, and three-times subqueries for the Sections 1 to 3, respectively. Bold values represent the best values in each row.

| Query approach | Federated or Centralized | Target diseases | The runtime of 1 subquery (for whole of sections) [x=1] | The runtime of 2 subqueries (for whole of sections) [x=2] | The runtime of 3 subqueries (for whole of sections) [x=3] | Mean | Standard deviation | The difference between the Federated and the Centralized means |
| --- | --- | --- | --- | --- | --- | --- | --- | --- |
| Example 1-x | Federated | AD | 59.3 | 47.5 | **46.6** | 51.3 | 7.1 | 24.8 |
| Example 2-x | Centralized |  | 40.0 | **11.6** | 27.9 | 26.5 | 14.3 |  |
| Example 3-x | Federated | melanoma | **38.1** | 41.7 | 40.2 | 40.5 | 1.8 | 6.6 |
| Example 4-x | Centralized |  | **19.6** | 42.0 | 40.2 | 33.9 | 12.4 |  |
|  |  | Mean | 39.3 | 35.7 | 38.7 |  |  |  |
|  |  | Standard deviation | 16.2 | 16.3 | 7.8 |  |  |  |

In all Examples, we arranged Section 4 to nest other Sections. Next, we measured the runtimes from Section 1 to 3 and that of Section 4 to presume the breakdown of the runtimes. Table 5 shows the runtimes of Sections 1 through 3. In both rows of “Example 5-x for AD (**Example 5-0** without subqueries (see Additional file 13), **5-1** with one subquery (see Additional file 14), and **5-2** with two subqueries (see Additional file 15)) and Example 6-x for melanoma (Example **6-0** without subqueries (see Additional file 16), **6-1** with one subquery (see Additional file 17), and **6-2** with two subqueries (see Additional file 18)), the runtimes of the queries with one or two subqueries (e.g., Example 5-1, 5-2, 6-1 and 6-2) were significantly faster than those without subqueries (e.g., Example 5-0 and 6-0) (Table 5). These results indicated that sections 1 through 3 took approximately 2 seconds to process. The retrieved bioresources and disease-related genes of Examples 5-0, 5-1, and 5-2 were the same, and those of 6-0, 6-1, and 6-2 were the same (Table 5).

**Table 5.**
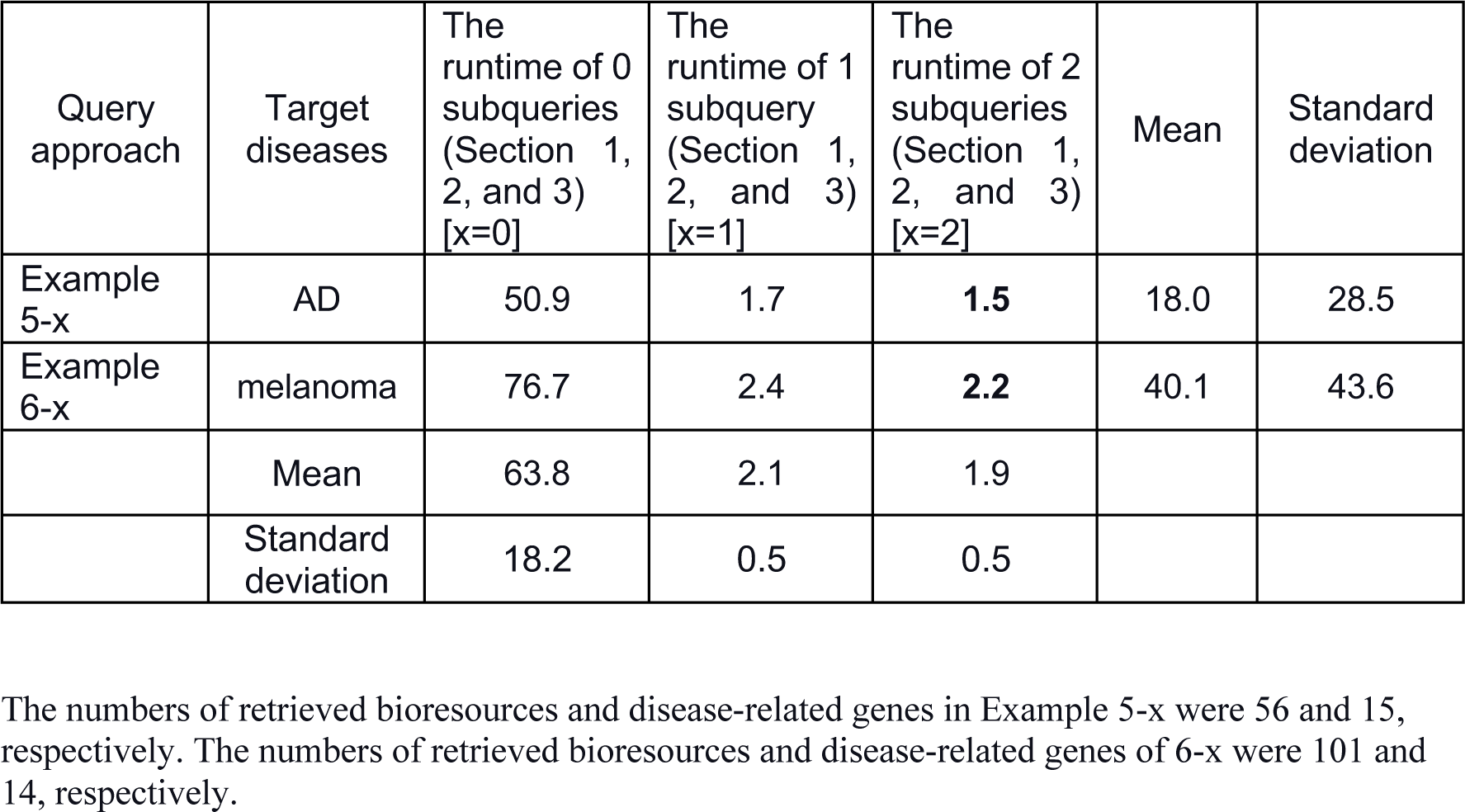
The average runtime from 10 executions of the SPARQL query Examples 5-x and 6-x, including one-time, twice, and three-times subqueries for the Sections 1 to 3, respectively. Bold values represent the best values in each row.

| Query approach | Target diseases | The runtime of 0 subqueries (Section 1, 2, and 3) [x=0] | The runtime of 1 subquery (Section 1, 2, and 3) [x=1] | The runtime of 2 subqueries (Section 1, 2, and 3) [x=2] | Mean | Standard deviation |
| --- | --- | --- | --- | --- | --- | --- |
| Example 5-x | AD | 50.9 | 1.7 | **1.5** | 18.0 | 28.5 |
| Example 6-x | melanoma | 76.7 | 2.4 | **2.2** | 40.1 | 43.6 |
|  | Mean | 63.8 | 2.1 | 1.9 |  |  |
|  | Standard deviation | 18.2 | 0.5 | 0.5 |  |  |

The numbers of retrieved bioresources and disease-related genes in Example 5-x were 56 and 15, respectively. The numbers of retrieved bioresources and disease-related genes of 6-x were 101 and 14, respectively.

Table 6 shows the runtime of Section 4. The runtimes of the federated search execution for the prefrontal cortex (**Example 7** (see Additional file 19)) and the skin of body (**Example 8** (see Additional file 20)) were both approximately 29 seconds, while those of the centralized search execution for the prefrontal cortex (**Example 9** (see Additional file 21)) and the skin of body (**Example 10** (see Additional file 22)) were approximately 9 and 11 seconds, respectively. The time differences between the federated and centralized approaches for AD and melanoma were approximately 20 and 18 seconds, respectively. The retrieved bioresources and disease-related genes were the same among Examples 7 and 9, and Examples 8 and 10, respectively (Table 6).

**Table 6.** The average runtime from 10 executions of the SPARQL query Examples 7, 8, 9, and 10 without using the subqueries in Section 4. Bold values represent the best values in each column. We executed the federated search from the BioResource MetaDB SPARQL endpoint to the official Bgee SPARQL endpoint. We performed the centralized search from the BioResource MetaDB SPARQL endpoint to the Bgee data stored in the BioResource MetaDB.

| Query approach | Federated or Centralized | Anatomical parts where genes were overexpressed | The runtime of 0 subqueries (Section 4) | The difference between the Federated and the Centralized |
| --- | --- | --- | --- | --- |
| Example7 | Federated | prefrontal cortex | 28.8 | 19.8 (prefrontal cortex), and 18.3 (skin of body) |
| Example8 |  | skin of body | 29.2 |  |
| Example9 | Centralized | prefrontal cortex | **9.0** |  |
| Example10 |  | skin of body | 10.9 |  |

The numbers of retrieved genes in Examples 7 and 9 were the same at 618. The numbers of retrieved genes in Examples 8 and 10 were the same and 498, respectively.

Moreover, we measured the runtime of the federated approach between the BioResource MetaDB (Tsukuba in Japan) and the Bgee (Lausanne in Switzerland), and the centralized approach for Bgee data in Tsukuba and Lausanne (Table 7). We executed the centralized approaches for Bgee data stored at the RIKEN BRC (Tsukuba) and the SIB (Lausanne), from each place. The executed query includes the query conditions: the prefrontal cortex (UBERON:0000451) as the location of overexpression, a high confidence level for expression data, and the expression score greater than 99. As a result, the runtime of the federated approach (Tsukuba to Lausanne) was approximately 29 seconds, including data transfer time and the Bgee triple store query evaluation time. The centralized approach runtime in Lausanne (Lausanne to Lausanne) was about 26 seconds, and that in Tsukuba (Tsukuba to Tsukuba) was approximately 9 seconds. From these results, we estimated the data transfer time between Tsukuba and Lausanne was approximately 2-3 seconds, and the difference between the query evaluation time of the BioResource MetaDB in Tsukuba and Bgee in Lausanne was approximately 17 seconds.

**Table 7.** Comparison of the runtimes of the federated approach from the BioResource MetaDB (Tsukuba) to the Bgee (Lausanne), and the centralized approach at Tsukuba and Lausanne. We conducted the performance by executing the SPARQL query for Bgee data (Examples 7 and 9).

| Query approach | Mean (seconds) | Standard deviation (seconds) | Results (cases) | A-B (seconds) | B-C (seconds) |
| --- | --- | --- | --- | --- | --- |
| Federated (Tsukuba to Lausanne): Example 7 | 28.8 [A] | 0.7 | 618 | 2.4 |  |
| Centralized (Lausanne to Lausanne) * | 26.4 [B] | 0.2 |  |  | 17.4 |
| Centralized (Tsukuba to Tsukuba): Example 9 | 9.0 [C] | 0.3 |  |  |  |

\*: After we removed a row including the SERVICE keyword from Example 7 and executed it.
A-B: Data transfer time between Tsukuba (RIKEN BRC) and Lausanne (Bgee)
B-C: The difference in the retrieval time of the BioResource MetaDB and Bgee

From the results of Tables 4, 5, 6, and 7, we concluded that one of the reasons for the query performance degradation in the federated approach and the improvement was as follows, (1) the difference in the total runtime of the federated and centralized approach (e.g., about 25 seconds in the case of AD in Table 4) mainly depended on the data transfer time between Tsukuba and Lausanne and the query evaluation time of Section 4 (Bgee data) since the runtime of Section 1 through 3 took approximately 2 seconds by using the subqueries (Table 5). (2) We estimated the data transfer time between Tsukuba and Lausanne took approximately 2 seconds (Table 7). At this time, the number of data transferred from Tsukuba to Lausanne was 56 cases (Table 5). We identified that the data transfer time was short since we refined the quantity of transferred data from Tsukuba to Lausanne using the subqueries. (3) On the other hand, the query evaluation times of Bgee data (Section 4) in the BioResource MetaDB (Tsukuba) and the Bgee database (Lausanne) took approximately 9 seconds and 26 seconds, respectively, and the difference between them was approximately 17 seconds (Table 7).

As a result, the reasons for the difference depend on the server’s specification (e.g., the memory capacity) and the database type (e.g., Virtuoso), the versions, the settings, and scalability issues. Therefore, we could improve the degradation of the query performance of the federated approach from the BioResource MetaDB to the SIB by enhancing the server specifications and by optimizing the triple store. First, we considered longer runtimes in the federated approach could be due to the network latency as one of the reasons (Section 6); however, the results suggested that the impact of network latency on federated search speed was insignificant, and its extent could be mitigated by reducing the quantity of data transfer during the execution of subqueries. Indeed, by optimizing the evaluation of triple store queries in Bgee’s triple store as an additional query performance test, the execution times of Examples 1-1 and 3-1 for the federated approach were improved to the same level as that of Examples 2-1 and 4-1 for the centralized approach. (see README.md in this project [48]).

In addition, using the federated search exhibited several important advantages. For institutions such as the RIKEN BRC, which combines its own RDF data with external datasets, using the federated approach should leverage the latest, most up-to-date information from each external dataset and thereby reduce operational costs that would be required to maintain a local copy in-sync when the external sources are updated. The federated approach is therefore particularly beneficial for institutions that use multiple third-party datasets. The federated approach is an essential technology for exploring bioresources relevant to biomedical research, which needs the combination of several external datasets.

### 7.2 Execution of a transitive search using external data

This study used Uberon ontology terms, such as prefrontal cortex (UBERON:0000451) or skin of body (UBERON:0002097), as anatomical parts where genes are expressed. However, we realized we could not comprehensively acquire overexpressed genes at specific anatomical locations using the Example queries shown so far. For example, in Examples 3-1 and 4-1, we specified the “skin of body” as the target anatomical parts and observed genes overexpressed at those anatomical parts. However, in these cases, we cannot find expressed genes on the “zone of skin” (UBERON:0000014) that is a part of “skin of body” or on the “skin of limb” (UBERON:0001419) that is subClassOf “zone of skin” (Fig. 6). When the users specify “skin of body” as target anatomical parts, they would often expect to acquire expression information from both the “skin of body” and the subclass concepts that are subClassOf or part of “skin of body.”

**Fig. 6.**
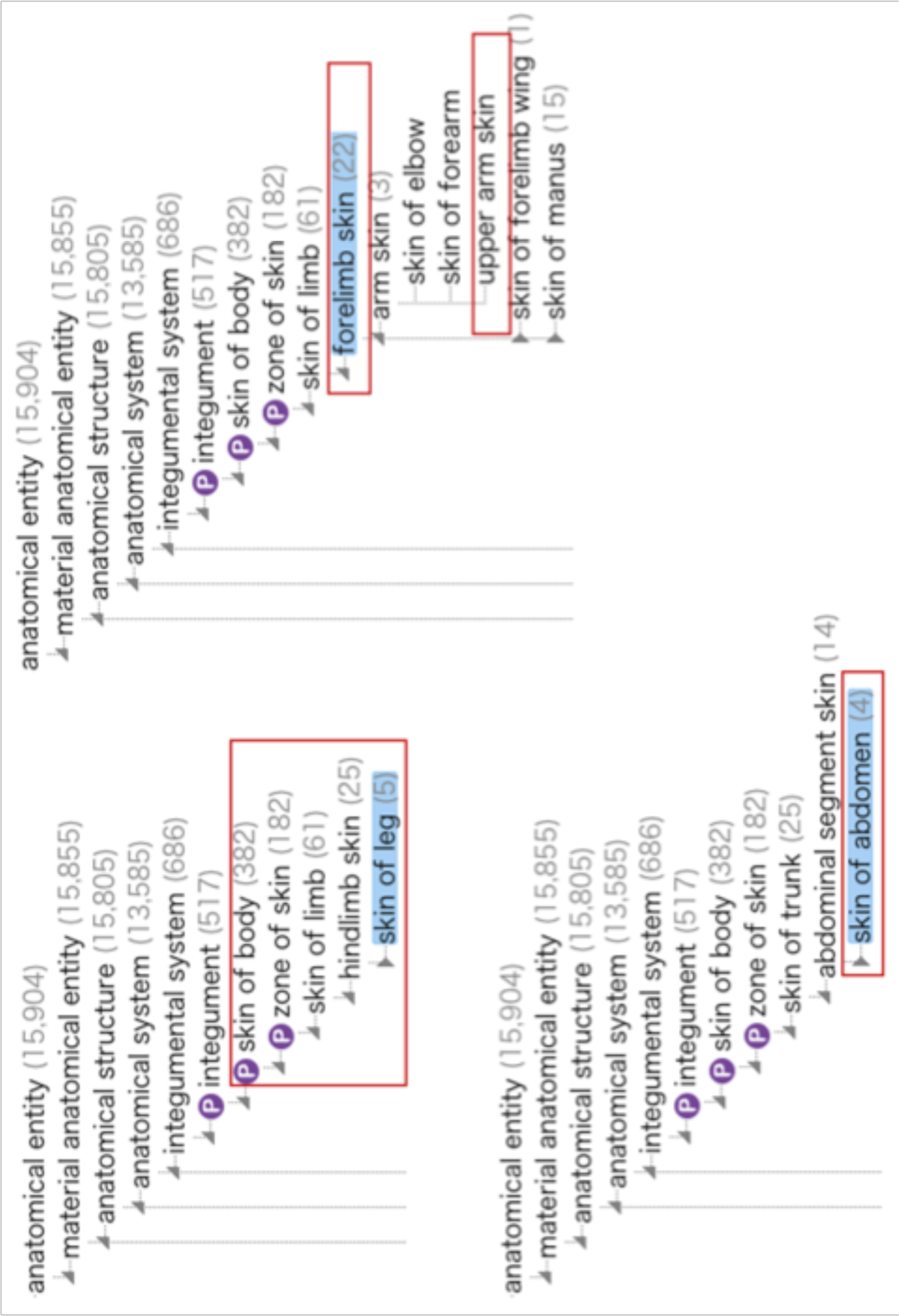
A part of the ontological tree of the Uberon ontology. Red rectangles indicate anatomical sites where melanoma-related genes were overexpressed. The “P” mark represents the “part of” relation. This ontological tree was made from a diagram of the Ontology Lookup Service (OLS) at https://www.ebi.ac.uk/ols4.

Balhoff et al. [24] cited “index_finger is_a finger” and “finger part_of_hand” as examples and they mentioned that a user would expect that when querying for parts of the hand they would receive not only ’finger’ but any concepts stated to be parts thereof (e.g., fingernails) or subclasses of ’finger,’ and SPARQL property paths cannot be easily employed to retrieve nodes linked by a chain of properties over such OWL expressions. Furthermore, OBO library ontologies include a wealth of inter-ontology semantic links, which require OWL reasoning to be fully utilized. One way to accomplish this would be to import all the needed ontologies into the Protégé tool [49, 50], and run an OWL reasoner, while it will need to be aware of the OWL-RDF serialization in order to match these complex triple patterns. Subsequently, they developed the Ubergraph, which currently includes 39 OBO ontologies including the Uberon with precomputed relations, to solve this issue by performing SPARQL queries that make use of the semantics of the included ontologies [24].

On the other hand, we strove to solve the problem of mixed subClassOf and partOf relationships between anatomical terms in Uberon, where the depth of the hierarchy is unknown, by reusing existing public resources and using SPARQL. We acquired the latest uberon_kgx_tsv_edge.tsv [51] that was published from the KG-OBO project and converted the downloaded tsv format file to two turtle (ttl) format files by a Python script (see Additional file 23). The uberon_kgx_tsv_edge.tsv was a KGX TSV format file by being transformed from uberon.owl [52] using the Koza tool [15]. Our converted two ttl format files included subject_broader_object_from_BFO_0000050.ttl (see Additional file 24) and subject_broader_object_from_subClassOf.ttl (see Additional file 25). The former file was converted from part of the relation between subject and object terms to the “broader” predicate [53], the latter file was converted from subClassOf relation to the “broader” predicate. The broader relation is a predicate directly connecting among uberon terms instead of partOf and subClassOf relations. We stored these two ttl format files as a named GRAPH: <http://metadb.riken.jp/db/uberonRDF_broader_fromKGX> into the BioResource MetaDB. We term these two ttl format data the uberonRDF-KGX.

Fig. 7 demonstrates a path between the “skin of limb” (UBERON:0001419) and the “skin of body” (UBERON:0002097) in the uberon.owl (diagram A) and the named GRAPH <http://metadb.riken.jp/db/uberonRDF_broader_fromKGX> (diagram B) within the RIKEN BioResource MetaDB. In the uberon.owl (diagram A), the “skin of body” connects to the “skin of limb” through the rdfs:subClassOf and owl:someValueFrom, while in the diagram B, the “skin of body” connects to the “skin of limb” through two broader predicates. Since it is difficult to execute a transitive search among Uberon terms by using the SPARQL query for uberon.owl (diagram A), we successfully executed a transitive search by using the Property Paths function of SPARQL query for the named GRAPH <http://metadb.riken.jp/db/uberonRDF_broader_fromKGX> (diagram B), whereby data was converted from part of and subClassOf relations to the broader predicate.

**Fig. 7.**
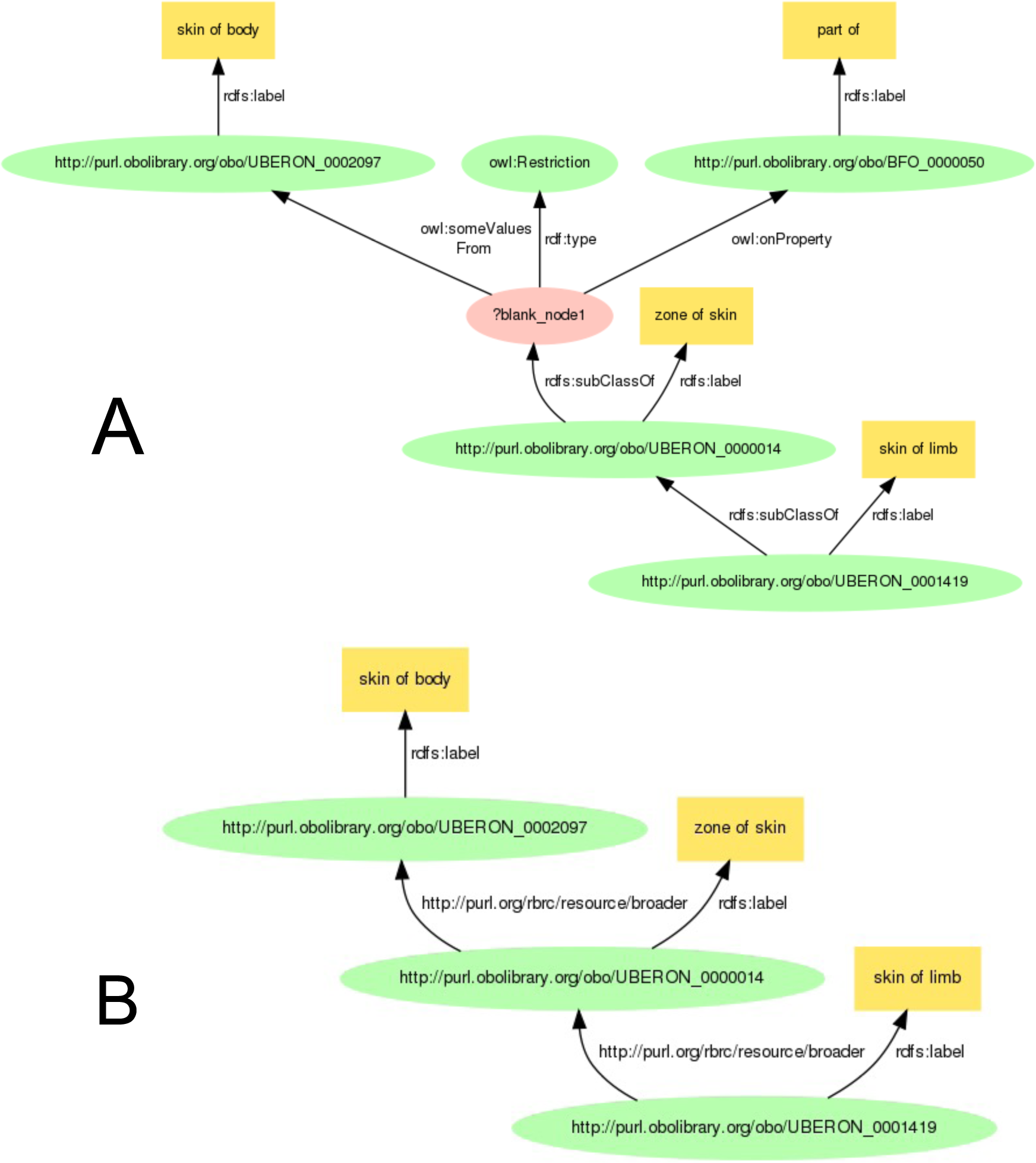
A path between the “skin of limb” (UBERON:0001419) and the “skin of body” (UBERON:0002097) in the uberon.owl (A) and that in the named GRAPH < http://metadb.riken.jp/db/uberonRDF_broader_fromKGX> within the RIKEN BioResource MetaDB (B). These diagrams were created using https://www.kanzaki.com/works/2009/pub/graph-draw.

**Example 11-1:** Centralized query for melanoma using the uberonRDF-KGX (see Additional file 26) is a SPARQL query where we added the named GRAPH: <http://metadb.riken.jp/db/uberonRDF_broader_fromKGX> to the Example 4-1 so as to execute a transitive search for the Uberon terms by using the Property Paths function.

**Example 11-2**: Centralized query for melanoma using the Ubergraph data instead of the uberonRDF-KGX is a SPARQL query (see Additional file 27). This query includes a service keyword to execute a transitive search for Uberon RDF data in the Ubergraph through the federated approach to the Ubergraph SPARQL endpoint [54]. In advance, we performed a preliminary test for Examples 11-1 and 11-2, identifying the same results.

Table 8 shows the average runtimes of Examples 11-1 and 11-2. The runtime of Example 11-1 was 67 seconds, on the other hand, we did not obtain the result of Example 11-2 due to the transaction timeout (over 10 minutes). Table 9 shows the query result of Example 11-1. We found the HRAS gene (ENSG:00000174775) and PTEN gene (ENSG:00000171862), which were overexpressed in the “skin of body” or seven anatomical locations that comprise the partOf or subClassOf the skin of body (Table 9). HRAS and PTEN genes are highly relevant for melanoma research, as shown in [55, 56]. The anatomical locations on which the HRAS and PTEN genes were overexpressed include eight locations, such as the skin of limb and forelimb skin (UBERON:0003531) in addition to the skin of body (Table 9, Fig. 6). Furthermore, we explored 21 RIKEN bioresources expected to be suitable for melanoma research (Table 9). The number of melanoma-related genes, anatomical parts characterized by overexpression, and bioresources predicted to be suitable for melanoma research in Example 11-1 (Table 9) were greater than those in Example 4-1 (Table 3). We concluded that this is because Example 11-1 could execute a transitive search for the Uberon data using the SPARQL query’s Property Paths function.

**Table 8.** The average runtime from 10 executions of the SPARQL query Examples 11-1 and 11-2.

| Query approach | Target diseases | Which Uberon data was used | The runtime (seconds) |
| --- | --- | --- | --- |
| Example11-1 | melanoma | The Uberon data converted from KGX stored in the BioResource MetaDB (Centralized approach) | 67 |
| Example11-2 |  | The Ubergraph data through the SPARQL endpoint (Federated approach) | Transaction timed out (over 10 mins) |

**Table 9.** Results of Example 11-1: Centralized query for melanoma using the broader predicate to perform the property path function.

| Query approach | No. of mice | RIKEN Mouse IDs | No. of genes | Ensemble gene IDs | Gene labels | No. of anatomical entities | Uberon IDs | Anatomical entity labels |
| --- | --- | --- | --- | --- | --- | --- | --- | --- |
| Example 11-1:<br>Centralized query for melanoma using the broader predicate | 21 | RBRC11578<br>RBRC02473<br>RBRC02481<br>RBRC02449<br>RBRC02488<br>RBRC02464<br>RBRC02485<br>RBRC02484<br>RBRC10866<br>RBRC02466<br>RBRC02479<br>RBRC02450<br>RBRC02586<br>RBRC02460<br>RBRC02451<br>RBRC02475<br>RBRC01088<br>RBRC02487<br>RBRC-AES1453<br>RBRC02300<br>RBRC02301 | 2 | ENSG00000174775<br>ENSG00000171862 | HRAS<br>PTEN | 8 | UBERON:0000014<br>UBERON:0001511<br>UBERON:0003532<br>UBERON:0001416<br>UBERON:0001419<br>UBERON:0002097<br>UBERON:0003531<br>UBERON:0004263 | zone of skin,<br>skin of leg,<br>hindlimb skin,<br>skin of abdomen,<br>skin of limb,<br>skin of body,<br>forelimb skin,<br>upper arm skin |

## 8. Future work

In the demonstration of Section 5, we only used the DisGeNET as a GDA dataset. However, in the preliminary trials we performed, we successfully demonstrated the use of other datasets, such as MedGen, and MGI, instead of the DisGeNET (see this project webpage [57]). Therefore, we can select one of these GDA datasets or combine several. In the latter case, we can use common (intersection of) GDA data among DisGeNET, MedGen, and MGI datasets. In addition, the integration of the Bgee dataset allows us to handle information on genes overexpressed or underexpressed at specific anatomical locations. The Bgee dataset includes the development stage (e.g., late adult stage), sex, strain, and data source (e.g., RNA Seq) in addition to the anatomical location. The use of Bgee gene expression data is expected to lead to the exploration of more specific disease-related genes and bioresources.

In this article, we introduced a method to explore bioresources used for specific disease research using SPARQL queries. However, not all users of bioresources can perform information retrieval using SPARQL. Furthermore, the SPARQL query’s runtime sometimes takes several tens of seconds depending on the query conditions (Tables 2 and 4), and we observed that it needs to be shorter to provide efficient retrieval results for users. Therefore, we have developed a keyword search engine and interface for bioresource users and have accomplished a few seconds of runtime. The Search for bioresources tab [2, 58] leverages the technology of SPARQList [59], which provides a REST API server for a SPARQL query against bioresource association data collected by crawling the KG (Fig. 3) [60] and is a bioresource search service that enables keyword search using disease name, gene name, resource name, and species name. We plan to expand the keyword search function in the Search for bioresources tab to enable searching by the Uberon Ontology term. Moreover, we are also developing an interface that allows users to select ontology terms from the ontology tree structure so as to search for the related bioresources.

## Abbreviations

AD: Alzheimer’s disease
BRC: BioResource Research Center
GDA: Gene Disease Associations
KG: Knowledge Graphs
MGI: Mouse Genome Informatics
OMA: Orthologous MAtrix
RDF: Resource Description Framework

## Additional file 1

\# Query Example 1-1

\# Example 1-1: A federated SPARQL query for Alzheimer’s disease using one subquery

\# Example1-1_Additional_file_1.txt

\# https://github.com/kushidat/broaderPredicate_uberon/blob/main/Example1-1_Additional_file_1.txt

## Additional file 2

\# Query Example 2-1

\# Example 2-1: A centralized SPARQL query for Alzheimer’s disease with one subquery

\# Example2-1_Additional_file_2.txt

\# https://github.com/kushidat/broaderPredicate_uberon/blob/main/Example2-1_Additional_file_2.txt

## Additional file 3

\# Query Example 3-1

\# Example 3-1: A federated SPARQL query for melanoma with one subquery

\# Example3-1_Additional_file_3.txt

\# https://github.com/kushidat/broaderPredicate_uberon/blob/main/Example3-1_Additional_file_3.txt

## Additional file 4

\# Query Example 4-1

\# Example 4-1: A centralized SPARQL query for melanoma with one subquery

\# Example4-1_ Additional_file_4.txt

\# https://github.com/kushidat/broaderPredicate_uberon/blob/main/Example4-1_Additional_file_4.txt

## Additional file 5

\# Query Example 1-2

\# Example 1-2: A federated SPARQL query for Alzheimer’s disease with two subqueries

\# Example1-2_ Additional_file_6.txt

\# https://github.com/kushidat/broaderPredicate_uberon/blob/main/Example1-2_Additional_file_5.txt

## Additional file 6

\# Query Example 2-2

\# Example 2-2: A centralized SPARQL query for Alzheimer’s disease with two subqueries

\# Example2-2_ Additional_file_6.txt

\# https://github.com/kushidat/broaderPredicate_uberon/blob/main/Example2-2_Additional_file_6.txt

## Additional file 7

\# Query Example 3-2

\# Example 3-2: A federated SPARQL query for melanoma with two subqueries

\# Example3-2_ Additional_file_7.txt

\# https://github.com/kushidat/broaderPredicate_uberon/blob/main/Example3-2_Additional_file_7.txt

## Additional file 8

\# Query Example 4-2

\# Example 4-2: A centralized SPARQL query for melanoma with two subqueries

\# Example4-2_ Additional_file_8.txt

\# https://github.com/kushidat/broaderPredicate_uberon/blob/main/Example4-2_Additional_file_8.txt

## Additional file 9

\# Query Example 1-3

\# Example 1-3: A federated query for Alzheimer’s disease with three subqueries

\# Example1-3_Additional_file_98.txt

\# https://github.com/kushidat/broaderPredicate_uberon/blob/main/Example1-3_Additional_file_9.txt

## Additional file 10

\# Query Example 2-3

\# Example 2-3: A centralized SPARQL query for Alzheimer’s disease with three subqueries

\# Example2-3_ Additional_file_10.txt

\# https://github.com/kushidat/broaderPredicate_uberon/blob/main/Example2-3_Additional_file_10.txt

## Additional file 11

\# Query Example 3-3

\# Example 3-3: A federated SPARQL query for melanoma with three subqueries

\# Example3-3_ Additional_file_11.txt

\# https://github.com/kushidat/broaderPredicate_uberon/blob/main/Example3-3_Additional_file_11.txt

## Additional file 12

\# Query Example 4-3

\# Example 4-3: A centralized SPARQL query for melanoma with three subqueries

\# Example4-3_ Additional_file_12.txt

\# https://github.com/kushidat/broaderPredicate_uberon/blob/main/Example4-3_Additional_file_12.txt

## Additional file 13

\# Query Example 5-0

\# Example 5-0: A centralized SPARQL query of Sections 1 through 3 for Alzheimer’s disease without subqueries

\# Example5-0_ Additional_file_13.txt

\# https://github.com/kushidat/broaderPredicate_uberon/blob/main/Example5-0_Additional_file_13.txt

## Additional file 14

\# Query Example 5-1

\# Example 5-1: A centralized SPARQL query of Sections 1 through 3 for Alzheimer’s disease with one subquery

\# Example5-1_ Additional_file_14.txt

\# https://github.com/kushidat/broaderPredicate_uberon/blob/main/Example5-1_Additional_file_14.txt

## Additional file 15

\# Query Example 5-2

\# Example 5-2: A centralized SPARQL query of Sections 1 through 3 for Alzheimer’s disease with two subqueries

\# Example5-2_ Additional_file_15.txt

\# https://github.com/kushidat/broaderPredicate_uberon/blob/main/Example5-2_Additional_file_15.txt

## Additional file 16

\# Query Example 6-0

\# Example 6-0: A centralized SPARQL query of Sections 1 through 3 for melanoma without subqueries

\# Example6-0_ Additional_file_16.txt

\# https://github.com/kushidat/broaderPredicate_uberon/blob/main/Example6-0_Additional_file_16.txt

## Additional file 17

\# Query Example 6-1

\# Example 6-1: A centralized SPARQL query of Sections 1 through 3 for melanoma with one subquery

\# Example6-1_ Additional_file_17.txt

\# https://github.com/kushidat/broaderPredicate_uberon/blob/main/Example6-1_Additional_file_17.txt

## Additional file 18

\# Query Example 6-2

\# Example 6-2: A centralized SPARQL query of Sections 1 through 3 for melanoma with two subqueries

\# Example6-2_ Additional_file_18.txt

\# https://github.com/kushidat/broaderPredicate_uberon/blob/main/Example6-2_Additional_file_18.txt

## Additional file 19

\# Query Example 7

\# Example 7: A federated SPARQL query of Section 4 (Bgee) for the overexpressed genes in the “prefrontal cortex”

\# Example7_ Additional_file_19.txt

\# https://github.com/kushidat/broaderPredicate_uberon/blob/main/Example7_Additional_file_19.txt

## Additional file 20

\# Query Example 8

\# Example 4-1: A federated SPARQL query of Section 4 (Bgee) for the overexpressed genes in the “skin of body”

\# Example8_ Additional_file_20.txt

\# https://github.com/kushidat/broaderPredicate_uberon/blob/main/Example8_Additional_file_20.txt

## Additional file 21

\# Query Example 9

\# Example 4-1: A centralized SPARQL query of Section 4 (Bgee) for the overexpressed genes in the “prefrontal cortex”

\# Example9_ Additional_file_21.txt

\# https://github.com/kushidat/broaderPredicate_uberon/blob/main/Example9_Additional_file_21.txt

## Additional file 22

\# Query Example 10

\# Example 4-1: A centralized SPARQL query of Section 4 (Bgee) for the overexpressed genes in the “skin of body”

\# Example10_ Additional_file_22.txt

\# https://github.com/kushidat/broaderPredicate_uberon/blob/main/Example10_Additional_file_22.txt

## Additional file 23

\# tsv2rdf_uberonKgx_to_uberon_broader20230723.py

\# This is a Python script for converting the latest uberon_kgx_tsv_edge.tsv from the kg-uberon webpage in the KG-OBO project (https://kg-hub.berkeleybop.io/kg-obo/uberon/) to two ttl format files, including subject_broader_object_from_BFO_0000050.ttl (Additional file 24) and subject_broader_object_from_subClassOf.ttl (Additional file 25).

#https://github.com/kushidat/broaderPredicate_uberon/blob/main/tsv2rdf_uberonKgx_to_uberon_broader20230723.py

## Additional file 24

\# subject_broader_object_from_BFO_0000050.ttl

\# This is a turtle file converted from the latest uberon_kgx_tsv_edge.tsv in the kg-uberon webpage in the KG-OBO project (https://kg-hub.berkeleybop.io/kg-obo/uberon/). This file includes the broader predicate’s relationships among terms created from the BFO:0000050 (pratOf) relations in the uberon_kgx_tsv_edge.tsv in the kg-uberon.

#https://github.com/kushidat/broaderPredicate_uberon/blob/main/subject_broader_object_from_BFO_0000050.ttl

## Additional file 25

\# subject_broader_object_from_subClassOf.ttl

\# This is a turtle file converted from the latest uberon_kgx_tsv_edge.tsv in the kg-uberon webpage in the KG-OBO project (https://kg-hub.berkeleybop.io/kg-obo/uberon/). This file includes the broader predicate’s relationships among terms created from the rdfs:subClassOf relations in the uberon_kgx_tsv_edge.tsv in the kg-uberon.

#https://github.com/kushidat/broaderPredicate_uberon/blob/main/subject_broader_object_from_subClassOf.ttl

## Additional file 26

\# Query Example 11-1

\# Example 11-1: A centralized query for melanoma using uberonRDF-KGX

\# Example11-1_Additional_file_26.txt

\# https://github.com/kushidat/broaderPredicate_uberon/blob/main/Example11-1_Additional_file_26.txt

## Additional file 27

\# Query Example 11-2

\# Example 11-2: A centralized query for melanoma using the ubergraph

\# Example11-2_Additional_file_27.txt

\# https://github.com/kushidat/broaderPredicate_uberon/blob/main/Example11-2_Additional_file_27.txt

## Declarations

### Ethics approval and consent to participate

Not applicable.

### Consent for publication

Not applicable.

### Availability of data and materials

All materials and data of this study are available from the corresponding author on request.

### Competing interests

The authors declare no competing interests.

### Funding

Funding from State Secretariat for Education, Research and Innovation (SERI) via ETHZ grant BG 02-072020 and EU Horizon 2020 INODE grant 863410. This work was supported in part by ROIS-DS-JOINT (027RP2022) to T. Kushida.

### Authors’ contributions

TK contributed to the conceptualization, data collection, analysis, visualization, funding acquisition, and manuscript writing. TM contributed to the conceptualization, progress management, analysis, and manuscript writing. AS contributed to the conceptualization and manuscript writing. CD contributed to the conceptualization, funding acquisition, and supervision. HC contributed to the conceptualization, methodology, and manuscript writing. FB contributed to the conceptualization, data collection, funding acquisition, manuscript writing and supervision. HM contributed to the conceptualization, funding acquisition, manuscript writing, and supervision. All authors reviewed and approved the final manuscript.

## Acknowledgement

We thank Daiki Usuda and Masanobu Uchida for the Bioresource data preparation and the BRC server and triple store optimization.

## Authors’ information

TK, TM, AS, CD, HC, FB, HM

## Notes

### Competing Interest Statement

The authors have declared no competing interest.

## References

1. Kobayashi N, Kume S, Lenz K, Masuya H: Riken metadatabase: a database platform for health care and life sciences as a microcosm of linked open data cloud, International Journal on Semantic Web and Information Systems (IJSWIS) 14 (2018) 140–164.

2. Masuya H, Usuda D, Nakata H, Yuhara N, Kurihara K, Namiki Y, Iwase S, Takada T, Tanaka N, Suzuki K, Yamagata Y, Kobayashi N, Yoshiki A, Kushida T: Establishment and application of information resource of mutant mice in RIKEN BioResource Research Center. Lab Anim Res. 2021 Jan 18;37(1):6. doi: 10.1186/s42826-020-00068-8. PMID: 33455583; PMCID: PMC7811887.

3. Altenhoff AM, Train CM, Gilbert KJ, Mediratta I, Mendes de Farias T, Moi D, Nevers Y, Radoykova HS, Rossier V, Warwick Vesztrocy A, Glover NM, Dessimoz C: OMA orthology in 2021: website overhaul, conserved isoforms, ancestral gene order and more. Nucleic Acids Res. 2021 Jan 8;49(D1):D373–D379. doi: 10.1093/nar/gkaa1007. PMID: 33174605; PMCID: PMC7779010.

4. Piñero J, Saüch J, Sanz F, Furlong LI: The DisGeNET cytoscape app: Exploring and visualizing disease genomics data. Comput Struct Biotechnol J. 2021 May 11;19:2960–2967. doi: 10.1016/j.csbj.2021.05.015. PMID: 34136095; PMCID: PMC8163863.

5. Mondo Disease Ontology [http://obofoundry.org/ontology/mondo.html] Accessed on 27 July 2023

6. Schriml LM, Munro JB, Schor M, et al. The Human Disease Ontology 2022 update. Nucleic Acids Res. 2022;50(D1):D1255–D1261. doi:10.1093/nar/gkab1063

7. The Orphanet Rare Disease ontology (ORDO) [https://www.orphadata.com/ordo/] Accessed on 27 July 2023

8. The Nanbyo Disease Ontology (NANDO) [http://nanbyodata.jp/ontology/nando] Accessed on 27 July 2023

9. Kushida T, Usuda D, Takada T, Yamagata Y Masuya H: Ontology Integration for Discovering Bioresources Contributing to Medical Science Research [abstract] ICBO 2022: International Conference on Biomedical Ontology, 2022 [https://icbo-conference.github.io/icbo2022/papers/ICBO-2022_paper_1944.pdf, 2022]

10. Bastian FB, Roux J, Niknejad A, Comte A, Fonseca Costa SS, de Farias TM, Moretti S, Parmentier G, de Laval VR, Rosikiewicz M, Wollbrett J, Echchiki A, Escoriza A, Gharib WH, Gonzales-Porta M, Jarosz Y, Laurenczy B, Moret P, Person E, Roelli P, Sanjeev K, Seppey M, Robinson-Rechavi M: The Bgee suite: integrated curated expression atlas and comparative transcriptomics in animals. Nucleic Acids Res. 2021 Jan 8;49(D1):D831–D847. doi: 10.1093/nar/gkaa793. PMID: 33037820; PMCID: PMC7778977.

11. Fernández-Breis JT, Chiba H, Legaz-García Mdel C, Uchiyama I: The Orthology Ontology: development and applications. J Biomed Semantics. 2016 Jun 4;7(1):34. doi: 10.1186/s13326-016-0077-x. PMID: 27259657; PMCID: PMC4893294.

12. Mendes de Farias T, Chiba H, Fernández-Breis JT: Leveraging logical rules for efficacious representation of large orthology datasets. In Proceedings of the 10th International Semantic Web Applications and Tools for Healthcare and Life Sciences (SWAT4HCLS) Conference (Vol. 2042). 2017, April. CEUR-WS. https://ceur-ws.org/Vol-2042/paper36.pdf

13. Shefchek KA, Harris NL, Gargano M, Matentzoglu N, Unni D, Brush M, Keith D, Conlin T, Vasilevsky N, Zhang XA, Balhoff JP, Babb L, Bello SM, Blau H, Bradford Y, Carbon S, Carmody L, Chan LE, Cipriani V, Cuzick A, Della Rocca M, Dunn N, Essaid S, Fey P, Grove C, Gourdine JP, Hamosh A, Harris M, Helbig I, Hoatlin M, Joachimiak M, Jupp S, Lett KB, Lewis SE, McNamara C, Pendlington ZM, Pilgrim C, Putman T, Ravanmehr V, Reese J, Riggs E, Robb S, Roncaglia P, Seager J, Segerdell E, Similuk M, Storm AL, Thaxon C, Thessen A, Jacobsen JOB, McMurry JA, Groza T, Köhler S, Smedley D, Robinson PN, Mungall CJ, Haendel MA, Munoz-Torres MC, Osumi-Sutherland D: The Monarch Initiative in 2019: an integrative data and analytic platform connecting phenotypes to genotypes across species. Nucleic Acids Res. 2020 Jan 8;48(D1):D704–D715. doi: 10.1093/nar/gkz997. PMID: 31701156; PMCID: PMC7056945.

14. Ringwald M, Richardson JE, Baldarelli RM, Blake JA, Kadin JA, Smith C, Bult CJ: Mouse Genome Informatics (MGI): latest news from MGD and GXD. Mamm Genome. 2022 Mar;33(1):4–18. doi: 10.1007/s00335-021-09921-0. Epub 2021 Oct 26. PMID: 34698891; PMCID: PMC8913530.

15. Caufield JH, Putman T, Schaper K, Unni DR, Hegde H, Callahan TJ, Cappelletti L, Moxon SAT, Ravanmehr V, Carbon S, Chan LE, Cortes K, Shefchek KA, Elsarboukh G, Balhoff J, Fontana T, Matentzoglu N, Bruskiewich RM, Thessen AE, Harris NL, Munoz-Torres MC, Haendel MA, Robinson PN, Joachimiak MP, Mungall CJ, Reese JT: KG-Hub-building and exchanging biological knowledge graphs. Bioinformatics. 2023 July 1;39(7):btad418. doi: 10.1093/bioinformatics/btad418. PMID: 37389415; PMCID: PMC10336030.

16. KG-Hub webpage [http://kghub.org/] Accessed on 27 July 2023

17. Unni DR, Moxon SAT, Bada M, Brush M, Bruskiewich R, Caufield JH, Clemons PA, Dancik V, Dumontier M, Fecho K, Glusman G, Hadlock JJ, Harris NL, Joshi A, Putman T, Qin G, Ramsey SA, Shefchek KA, Solbrig H, Soman K, Thessen AE, Haendel MA, Bizon C, Mungall CJ; Biomedical Data Translator Consortium: Biolink Model: A universal schema for knowledge graphs in clinical, biomedical, and translational science. Clin Transl Sci. 2022 Aug;15(8):1848–1855. doi: 10.1111/cts.13302. Epub 2022 June 6. PMID: 36125173; PMCID: PMC9372416.

18. Reese JT, Unni D, Callahan TJ, Cappelletti L, Ravanmehr V, Carbon S, Shefchek KA, Good BM, Balhoff JP, Fontana T, Blau H, Matentzoglu N, Harris NL, Munoz-Torres MC, Haendel MA, Robinson PN, Joachimiak MP, Mungall CJ: KG-COVID-19: A Framework to Produce Customized Knowledge Graphs for COVID-19 Response. Patterns (N Y). 2021 Jan 8;2(1):100155. doi: 10.1016/j.patter.2020.100155. Epub 2020 Nov 9. PMID: 33196056; PMCID: PMC7649624.

19. KG-OBO webpage [http://kghub.org/kg_obo/] Accessed on 27 July 2023

20. Jackson R, Matentzoglu N, Overton JA, Vita R, Balhoff JP, Buttigieg PL, Carbon S, Courtot M, Diehl AD, Dooley DM, Duncan WD, Harris NL, Haendel MA, Lewis SE, Natale DA, Osumi-Sutherland D, Ruttenberg A, Schriml LM, Smith B, Stoeckert CJ Jr, Vasilevsky NA, Walls RL, Zheng J, Mungall CJ, Peters B. OBO Foundry in 2021: operationalizing open data principles to evaluate ontologies. Database (Oxford). 2021 Oct 26;2021:baab069. doi: 10.1093/database/baab069. PMID: 34697637; PMCID: PMC8546234.

21. Gene Ontology Consortium; Aleksander SA, Balhoff J, Carbon S, Cherry JM, Drabkin HJ, Ebert D, Feuermann M, Gaudet P, Harris NL, Hill DP, Lee R, Mi H, Moxon S, Mungall CJ, Muruganugan A, Mushayahama T, Sternberg PW, Thomas PD, Van Auken K, Ramsey J, Siegele DA, Chisholm RL, Fey P, Aspromonte MC, Nugnes MV, Quaglia F, Tosatto S, Giglio M, Nadendla S, Antonazzo G, Attrill H, Dos Santos G, Marygold S, Strelets V, Tabone CJ, Thurmond J, Zhou P, Ahmed SH, Asanitthong P, Luna Buitrago D, Erdol MN, Gage MC, Ali Kadhum M, Li KYC, Long M, Michalak A, Pesala A, Pritazahra A, Saverimuttu SCC, Su R, Thurlow KE, Lovering RC, Logie C, Oliferenko S, Blake J, Christie K, Corbani L, Dolan ME, Drabkin HJ, Hill DP, Ni L, Sitnikov D, Smith C, Cuzick A, Seager J, Cooper L, Elser J, Jaiswal P, Gupta P, Jaiswal P, Naithani S, Lera-Ramirez M, Rutherford K, Wood V, De Pons JL, Dwinell MR, Hayman GT, Kaldunski ML, Kwitek AE, Laulederkind SJF, Tutaj MA, Vedi M, Wang SJ, D’Eustachio P, Aimo L, Axelsen K, Bridge A, Hyka-Nouspikel N, Morgat A, Aleksander SA, Cherry JM, Engel SR, Karra K, Miyasato SR, Nash RS, Skrzypek MS, Weng S, Wong ED, Bakker E, Berardini TZ, Reiser L, Auchincloss A, Axelsen K, Argoud-Puy G, Blatter MC, Boutet E, Breuza L, Bridge A, Casals-Casas C, Coudert E, Estreicher A, Livia Famiglietti M, Feuermann M, Gos A, Gruaz-Gumowski N, Hulo C, Hyka-Nouspikel N, Jungo F, Le Mercier P, Lieberherr D, Masson P, Morgat A, Pedruzzi I, Pourcel L, Poux S, Rivoire C, Sundaram S, Bateman A, Bowler-Barnett E, Bye-A-Jee H, Denny P, Ignatchenko A, Ishtiaq R, Lock A, Lussi Y, Magrane M, Martin MJ, Orchard S, Raposo P, Speretta E, Tyagi N, Warner K, Zaru R, Diehl AD, Lee R, Chan J, Diamantakis S, Raciti D, Zarowiecki M, Fisher M, James-Zorn C, Ponferrada V, Zorn A, Ramachandran S, Ruzicka L, Westerfield M: The Gene Ontology knowledgebase in 2023. Genetics. 2023 May 4;224(1):iyad031. doi: 10.1093/genetics/iyad031. PMID: 36866529; PMCID: PMC10158837.

22. Hastings J, Owen G, Dekker A, Ennis M, Kale N, Muthukrishnan V, Turner S, Swainston N, Mendes P, Steinbeck C. ChEBI in 2016: Improved services and an expanding collection of metabolites. Nucleic Acids Res. 2016 Jan 4;44(D1):D1214–9. doi: 10.1093/nar/gkv1031. Epub 2015 Oct 13. PMID: 26467479; PMCID: PMC4702775.

23. Mungall CJ, Torniai C, Gkoutos GV, Lewis SE, Haendel MA.: Uberon, an integrative multi-species anatomy ontology. Genome Biol. 2012;13(1):R5. Published 2012 Jan 31. doi:10.1186/gb-2012-13-1-r5

24. Balhoff JP, Bayindir U, Caron AR, Matentzoglu N, OsumiSutherland D Mungall CJ: Ubergraph: integrating OBO ontologies into a unified semantic graph. ICBO 2022: International Conference on Biomedical Ontology (ICBO) 2022, [https://icbo-conference.github.io/icbo2022/papers/ICBO-2022_paper_5005.pdf]

25. OWL 2 Web Ontology Language Document Overview (Second Edition) [https://www.w3.org/TR/owl2-overview/#sec-ont] Accessed on 27 July 2023

26. Diehl AD, Meehan TF, Bradford YM, Brush MH, Dahdul WM, Dougall DS, He Y, Osumi-Sutherland D, Ruttenberg A, Sarntivijai S, Van Slyke CE, Vasilevsky NA, Haendel MA, Blake JA, Mungall CJ: The Cell Ontology 2016: enhanced content, modularization, and ontology interoperability. J Biomed Semantics. 2016 July 4;7(1):44. doi: 10.1186/s13326-016-0088-7. PMID: 27377652; PMCID: PMC4932724.

27. Smith CL, Eppig JT. The mammalian phenotype ontology: enabling robust annotation and comparative analysis. Wiley Interdiscip Rev Syst Biol Med. 2009 Nov-Dec;1(3):390–399. doi: 10.1002/wsbm.44. PMID: 20052305; PMCID: PMC2801442.

28. Köhler S, Gargano M, Matentzoglu N, Carmody LC, Lewis-Smith D, Vasilevsky NA, Danis D, Balagura G, Baynam G, Brower AM, Callahan TJ, Chute CG, Est JL, Galer PD, Ganesan S, Griese M, Haimel M, Pazmandi J, Hanauer M, Harris NL, Hartnett MJ, Hastreiter M, Hauck F, He Y, Jeske T, Kearney H, Kindle G, Klein C, Knoflach K, Krause R, Lagorce D, McMurry JA, Miller JA, Munoz-Torres MC, Peters RL, Rapp CK, Rath AM, Rind SA, Rosenberg AZ, Segal MM, Seidel MG, Smedley D, Talmy T, Thomas Y, Wiafe SA, Xian J, Yüksel Z, Helbig I, Mungall CJ, Haendel MA, Robinson PN: The Human Phenotype Ontology in 2021. Nucleic Acids Res. 2021 Jan 8;49(D1):D1207-D1217. doi: 10.1093/nar/gkaa1043. PMID: 33264411; PMCID: PMC7778952.

29. RIKEN BRC webpage [https://web.brc.riken.jp/] Accessed on 27 July 2023

30. BRSO webpage [hhttps://github.com/dbcls/brso] Accessed on 27 July 2023

31. BioResource MetaDatabase webpage [https://knowledge.brc.riken.jp/sparql] Accessed on 27 July 2023

32. MGI_EntrezGene.rpt [http://www.informatics.jax.org/downloads/reports/MGI_EntrezGene.rpt] Accessed on 27 July 2023

33. UniProt ID idmapping_selected.tab.gz [https://ftp.uniprot.org/pub/databases/uniprot/current_release/knowledgebase/idmapping/idmapping_selected.tab.gz] Accessed on 27 July 2023

34. MedGen [https://www.ncbi.nlm.nih.gov/medgen/] Accessed on 27 July 2023

35. Kawashima S, Katayama T, Hatanaka H, Kushida T, Takagi T. NBDC RDF portal: a comprehensive repository for semantic data in life sciences. Database (Oxford). 2018 Jan 1;2018:bay123. doi: 10.1093/database/bay123. PMID: 30576482; PMCID: PMC6301334.

36. DisGeNET v.7.0.0 RDF data [https://rdfportal.org/download/disgenet/latest] Accessed on 27 July 2023

37. ICD-11 webpage [https://icd.who.int/] Accessed on 27 July 2023

38. Mendes de Farias T, Kushida T, Sima AC, Dessimoz C, Chiba H, Bastian F, Masuya H: Data in use for Alzheimer disease study: combining gene expression, orthology, bioresource and disease datasets. 14th International Conference on Semantic Web Applications and Tools for Health Care and Life Sciences (SWAT4HCLS 2023) 2023, 177–178. https://ceur-ws.org/Vol-3415/paper-47.pdf

39. BioResource MetaDatabase SPARQL endpoint [https://knowledge.brc.riken.jp/sparql] Accessed on 27 July 2023

40. Zheng H, Koo EH: The amyloid precursor protein: beyond amyloid. Mol Neurodegener. 2006 Jul 3;1:5. doi: 10.1186/1750-1326-1-5. PMID: 16930452; PMCID: PMC1538601.

41. Zalocusky KA, Najm R, Taubes AL, Hao Y, Yoon SY, Koutsodendris N, Nelson MR, Rao A, Bennett DA, Bant J, Amornkul DJ, Xu Q, An A, Cisne-Thomson O, Huang Y: Neuronal ApoE upregulates MHC-I expression to drive selective neurodegeneration in Alzheimer’s disease. Nat Neurosci. 2021 Jun;24(6):786–798. doi: 10.1038/s41593-021-00851-3. Epub 2021 May 6. PMID: 33958804; PMCID: PMC9145692.

42. Bgee RDF data [https://www.bgee.org/ftp/current/rdf_easybgee.zip] Accessed on 27 July 2023

43. RIKEN BRC Mouse RBRC10866 webpage [https://knowledge.brc.riken.jp/resource/animal/card?brc_no=RBRC10866&lang=en] Accessed on 27 July 2023

44. RIKEN BRC Mouse RBRC01088 webpage [https://knowledge.brc.riken.jp/resource/animal/card?brc_no=RBRC01088&lang=en] Accessed on 27 July 2023

45. Bgee SPARQL endpoint [https://bgee.org/sparql/] Accessed on 27 July 2023

46. Qudus U, Saleem M, Ngonga Ngomo AC, Lee Young-Kooa.: An Empirical Evaluation of Cost-based Federated SPARQL Query Processing Engines. 1 Jan. 2021 : 843–868. doi:10.3233/SW-200420

47. SPARQL 1.1 Query Language. W3C Recommendation 21 March 2013 [https://www.w3.org/TR/2013/REC-sparql11-query-20130321/] Accessed on 27 July 2023

48. README.md in this project webpage [https://github.com/kushidat/broaderPredicate_uberon/blob/main/README.md] Accessed on 27 July 2023

49. Musen MA, Protégé Team: The Protégé Project: A Look Back and a Look Forward. AI Matters. 2015 Jun;1(4):4–12. doi: 10.1145/2757001.2757003. PMID: 27239556; PMCID: PMC4883684.

50. Protégé webpage [https://protege.stanford.edu/] Accessed on 27 July 2023

51. uberon_kgx_tsv_edge.tsv [https://kg-hub.berkeleybop.io/kg-obo/uberon/] Accessed on 27 July 2023

52. uberon.owl [http://purl.obolibrary.org/obo/uberon.owl] Accessed on 27 July 2023

53. rbrc:broader predicate URI [http://purl.org/rbrc/resource/broader] Accessed on 27 July 2023

54. Ubergraph SPARQL endpoint [https://yasgui.triply.cc/#] Accessed on 27 July 2023

55. Nogueira C, Kim KH, Sung H, Paraiso KH, Dannenberg JH, Bosenberg M, Chin L, Kim M: Cooperative interactions of PTEN deficiency and RAS activation in melanoma metastasis. Oncogene. 2010 Nov 25;29(47):6222–32. doi: 10.1038/onc.2010.349. Epub 2010 Aug 16. PMID: 20711233; PMCID: PMC2989338.

56. Tsao H, Zhang X, Fowlkes K, Haluska FG: Relative reciprocity of NRAS and PTEN/MMAC1 alterations in cutaneous melanoma cell lines. Cancer Res. 2000 Apr 1;60(7):1800–4. PMID: 10766161.

57. This project webpage [https://github.com/kushidat/broaderPredicate_uberon/tree/main] Accessed on 27 July 2023

58. The Search for bioresources tab webpage [https://web.brc.riken.jp/] Accessed on 27 July 2023

59. Katayama T, Kawashima S: SPARQList: Markdown-Based Highly Configurable REST API Hosting Server for SPARQL. In Proceedings of the 10th International Conference on Semantic Web Applications and Tools for Health Care and Life Sciences (SWAT4LS 2017) 2017 [https://ceur-ws.org/Vol-2042/paper47.pdf]

60. BRC SPARQList webpage [https://splist.brc.riken.jp/sparqlist/] Accessed on 27 July 2023

